# Lectins and polysaccharide EPS I have flow-responsive roles in the attachment and biofilm mechanics of plant pathogenic *Ralstonia*

**DOI:** 10.1101/2024.06.21.599993

**Authors:** Mariama D. Carter, Tuan M. Tran, Matthew L. Cope-Arguello, Sofia Weinstein, Hanlei Li, Connor Hendrich, Jessica L. Prom, Jiayu Li, Lan Thanh Chu, Loan Bui, Harishankar Manikantan, Tiffany Lowe-Power, Caitilyn Allen

**Affiliations:** Department of Plant Pathology, University of Wisconsin-Madison, Madison, Wisconsin, United States of America; Department of Biology, University of South Alabama, Mobile, Alabama, United States of America; Department of Plant Pathology, University of California-Davis, Davis, California, United States of America; Department of Chemical Engineering, University of California-Davis, Davis, California, United States of America; Department of Biology, University of Dayton, Dayton, Ohio, United States of America

## Abstract

Bacterial biofilm formation and attachment to hosts are mediated by carbohydrate-binding lectins, exopolysaccharides, and their interactions in the extracellular matrix (ECM). During tomato infection *Ralstonia pseudosolanacearum* (*Rps*) GMI1000 highly expresses three lectins: LecM, LecF, and LecX. The latter two are uncharacterized. We evaluated the roles in bacterial wilt disease of LecF, a fucose-binding lectin, LecX, a xylose-binding lectin, and the *Rps* exopolysaccharide EPS I. Interestingly, single and double lectin mutants attached to tomato roots better and formed more biofilm under static conditions *in vitro*. Consistent with this finding, static bacterial aggregation was suppressed by heterologous expression of *lecF*_GMI1000_ and *lecX*_GMI1000_ in other *Ralstonia* strains that naturally lack these lectins. Crude ECM from a Δ*lecF/X* double mutant was more adhesive than the wild-type ECM, and LecF and LecX increased *Rps* attachment to ECM. The enhanced adhesiveness of the Δ*lecF/X* ECM could explain the double mutant’s hyper-attachment in static conditions. Unexpectedly, mutating lectins decreased *Rps* attachment and biofilm viscosity under shear stress, which this pathogen experiences in plant xylem. LecF, LecX, and EPS I were all essential for biofilm development in xylem fluid flowing through cellulose-coated microfluidic channels. These results suggest that under shear stress, LecF and LecX increase *Rps* attachment by interacting with the ECM and plant cell wall components like cellulose. In static conditions such as on root surfaces and in clogged xylem vessels, the same lectins suppress attachment to facilitate pathogen dispersal. Thus, *Rps* lectins have a dual biological function that depends on the physical environment.

**Author Summary:** Bacterial wilt diseases caused by *Ralstonia* species inflict significant losses on diverse, globally important agricultural plants. The pathogen first colonizes roots and ultimately the water-transporting xylem. There it attaches to host cell walls and other bacterial cells to form biofilms that eventually block xylem vessels and disrupt sap flow. It is not well known how *Ralstonia* spp. modulate attachment, but precise control of both attachment and dispersal is critical for successful host colonization over the disease cycle. Excessive adhesion could trap bacteria in a toxic or nutrient-depleted environment. Conversely, insufficient adhesion in a flowing environment could displace bacteria from an optimal niche. We provide evidence of dual, environment-specific roles of carbohydrate-binding lectins and exopolysaccharide EPS I in *Ralstonia pseudosolanacearum* (*Rps*) attachment. In static conditions, which *Rps* experiences on a host root, two lectins suppress bacterial aggregation and adhesion to roots. However, in flowing conditions, which *Rps* experiences in healthy xylem vessels, the same two lectins and EPS I are essential for biofilm development. The lectins increase the biofilm viscosity and support colony structural integrity, likely by interacting with polysaccharides in the biofilm matrix. This novel multifunctionality of bacterial lectins reveals how pathogens adapt to a physically dynamic host environment.

## Introduction

Bacterial pathogens often depend on attachment to a host, adhesion, and attachment to other microbial cells, cohesion. Together, these behaviors yield biofilms, which are aggregated microbes encased in a matrix of extracellular polymeric substances that includes extracellular polysaccharides (EPS), extracellular nucleic acid (eDNA and eRNA), membrane vesicles, lipids, and proteins [1,2]. This flexible and dynamic extracellular matrix (ECM) provides protection from environmental stresses, an external digestion system, resource capture, and also mediates social interactions between bacterial cells [1,3]. The structural stability of the ECM is particularly important in environments with physical perturbations like flow, which microbes can sense via rheosensing [4,5]. Bacteria must precisely regulate their attachment to optimize transitions between motile and sessile lifestyles. This regulation is especially critical for plant vascular wilt pathogens.

Bacterial wilt diseases of diverse plants are caused by strains in the *Ralstonia solanacearum* species complex (RSSC), three species that fall into four phylogenetically and geographically distinct phylotypes: *R. pseudosolanacearum* (*Rps*) includes phylotypes I and III from Asia and Africa, respectively; *R. solanacearum* (*Rs*) contains phylotypes IIA, IIB, and IIC from the Americas; and *R. syzygii* (*Rsy*) consists of phylotype IV from the Indonesian archipelago and Japan [6]. All RSSC colonize the water-transporting xylem vessels of plants, and most are soil-borne and infect plant hosts at the root [7]. Attracted by root exudates and increasing intracellular energy levels, these pathogens use flagella-mediate chemotaxis and aerotaxis to locate host roots [8,9]. Once at the root, they adhere to the rhizoplane using secreted polysaccharides, pili, and afimbrial adhesive proteins [10–13]. The pathogen exploits root wounds to colonize intercellular spaces of the root cortex, where it forms biofilms [14–16]. From there *Ralstonia* cells enter the root xylem and spread systemically, growing to very high cell density, and forming biofilms that occlude vascular flow and cause characteristic wilting symptoms [17]. Repeated cycles of attachment and dispersal are essential as this bacterium transitions from the rhizoplane to the xylem. What molecular mechanisms drive adhesion and cohesion in these physically and chemically diverse host environments?

Lectins, proteins that recognize and reversibly bind carbohydrates, are potential players in this process. Bacterial lectins mediate host-microbe interactions and biofilm formation [18–20]. The opportunistic animal and plant pathogen *Pseudomonas aeruginosa* uses glycan-binding lectins LecA and LecB for host attachment and cell invasion [21]. Both these lectins promote biofilm formation *in vitro* [22,23]. In addition to host glycans, some lectins can recognize self-produced exopolysaccharides. LecB, which resembles the RSSC lectin LecM, binds the polysaccharide Psl and coordinates proper localization of Psl within the EPS matrix [24]. The rice pathogen *Xanthomonas oryzae* pv. *oryzae* has an adhesin, XadM, that contributes to leaf attachment, entry, and virulence [25]. XadM aids in biofilm formation *in vitro* and mediates attachment to exopolysaccharides. In both *X. oryzae* and *P. aeruginosa*, extracellular polysaccharides play a critical role in biofilms and virulence [26,27]. Lectin-exopolysaccharide interactions are thus a broadly adapted strategy for biofilm formation.

In the RSSC, as in many other bacterial pathogens, attachment is regulated by a quorum sensing (QS) system via the global regulator PhcA, which is activated by the accumulation of the QS signal [10,11,13,28–32]. As *Ralstonia* populations grow in confined spaces, signal concentration increases, upregulating biosynthesis of exopolysaccharide EPS I and secreted enzymes that directly increase virulence and help bacteria disperse from biofilms [30,31]. EPS is a key bacterial wilt virulence factor and a major component of the bacterial biomass in vascular systems of infected plants [33]. EPS I-deficient mutants are severely impaired in wilt symptom development, stem colonization, and systemic spread in the host stem, but the direct function of EPS in RSSC biofilm formation is unclear [34–36]. Deleting the QS regulator PhcA, which locks the bacterium in a low cell density mode, results in a hyper-attachment phenotype in tomato plants and dysregulates 14 putative adhesin genes. We found that three adhesins upregulated in a Δ*phcA* strain, *rcpA*, *rcpB*, and *radA*, help *Rps* phylotype I sequevar 18 strain GMI1000 attach to roots [10]. In contrast, genes encoding lectin adhesins LecF, LecM, and LecX, which are among the pathogen’s most highly expressed genes *in vitro* and *in planta,* are strongly repressed in the Δ*phcA* mutant [37–40]. *Rs* LecM contributes to biofilm formation, virulence, and host colonization at cool temperatures in the *Rs* phylotype IIB-1 strain UW551 [41]. In *Rps* phylotype I strain OE1-1, LecM aids in biofilm formation *in vitro* and colonization of tomato roots and stems [42]. An OE1-1 *lecM* mutant was also avirulent on wilt-susceptible tomatoes, suggesting that LecM is a critical virulence factor in both phylotype I and II strains.

While *lecM* has been characterized in two RSSC strains, nothing is known about the biological roles of *lecF* and *lecX* in bacterial wilt. Purified LecF (also named RSL), a homotrimer with a six-bladed beta-propeller architecture, has high binding affinities for L-fucose, L-galactose, D-fructose, and fucose-containing oligosaccharides [37–39]. Consistent with a biological role in adhesion to plants, LecF also binds xyloglucan, a plant cell wall hemicellulose with D-xylose side chains that can also contain L-fucose and D-galactose [37,43]. LecX (also named RS20L), a homotrimer, has a high binding affinity for L-fucose and, to a lesser extent, D-mannose and has weak interactions with N-acetyl-D-glucosamine-containing oligosaccharides [44]. Some of these sugars are components of *Rps* exopolysaccharides, lipopolysaccharides, and surface exposed features [45,46]. The coordinated upregulation of lectins and EPS I at high cell densities in the stem suggests they might function synergistically.

LecF and LecX could bind EPS sugars in the ECM and thus support bacterial cohesion and biofilm stability; they might also bind glycans on plant cell walls and thus aid bacterial adhesion and biofilm establishment.

Working with targeted mutants in several RSSC strains, we used a suite of biochemical, *in vitro*, and *in planta* attachment assays to explore the roles of bacterial lectins and exopolysaccharides in the ECM, biofilm formation, and bacterial interactions with the tomato host plant. We found that both LecF and LecX reduce attachment of plant pathogenic *Ralstonia* under static conditions, such as on root surfaces. However, in an environment under flow such as in xylem vessels, LecF and LecX help *Rps* attach to surfaces and form biofilms. Further, deletion of *lecF* and *lecX* altered the physical properties of the ECM and bacterial attachment to the ECM, demonstrating that matrix components function as an interdependent system rather than in isolation.

## Results

### Lectins repress *Rps* attachment to tomato roots

When deletion of the *phcA* QS gene locks GMI1000 in a low cell density mode, expression of lectin genes *lecF*, *lecM*, and *lecX* is strongly repressed *in planta* [29]. This led us to hypothesize that *Rps* lectin gene expression is low in the earliest stage of disease at the host root surface, but then increases when the bacteria invade roots and reach high cell density in the stem. To test this, we used qRT-PCR to measure lectin gene expression in bacteria colonizing three distinct environments in the tomato host: cells attached to the rhizoplane at low cell density, cells colonizing the root endosphere, and cells colonizing and forming biofilms in the stem. Two control genes indicative of *Rps* cell density state, *iolG* and *epsB,* showed the expected expression patterns in these samples (Fig 1A). IolG, a myo-inositol catabolism enzyme, is upregulated at low cell density and is required for *Rps* root colonization [10,47]. Consistent with this, relative to cells on the root surface *iolG* expression trended higher in the root endosphere, but not in the stem. EpsB, a predicted tyrosine-protein kinase likely involved in EPS I secretion, is upregulated at high cell density and is required for EPS I production [10,30,33]. As expected, *epsB* expression increased in bacteria colonizing the endosphere and stem relative to the root surface.

**Figure 1.**
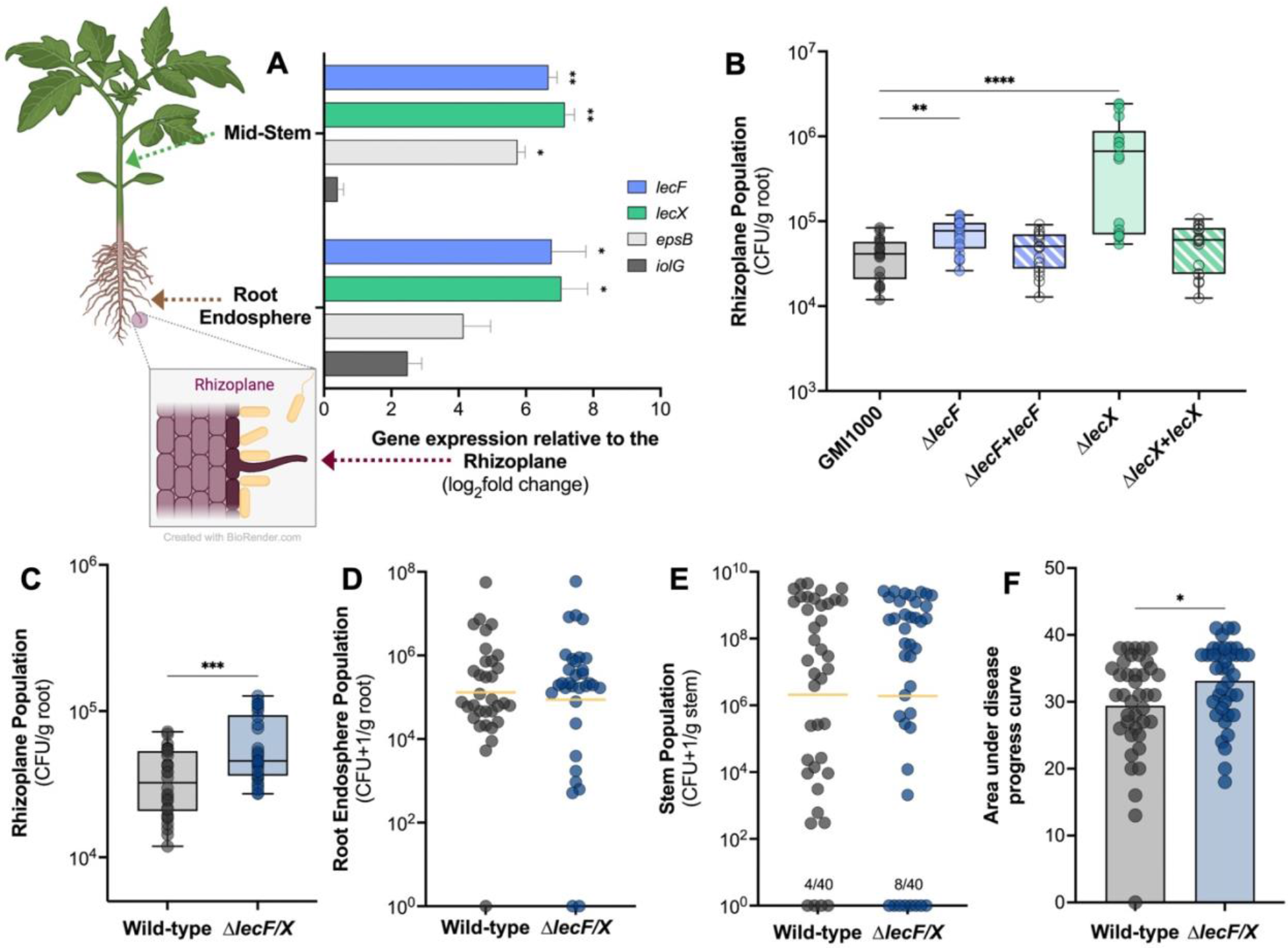
*In planta* expression and phenotypic contributions of *Ralstonia* lectin genes *lecF* and *lecX*. A) Expression of lectin genes *lecF* and *lecX* was upregulated in tomato root endosphere and stem. Five-day-old axenic tomato seedlings were flood-inoculated with a low cell density of *Rps* GMI1000. Rhizoplane and root endosphere samples were harvested 6 hpi and 48 hpi, respectively. Mid-stem samples were harvested from 24-day-old tomato plants three days after petiole inoculation. Total RNA was extracted from plant tissue samples and qRT-PCR was used to quantify expression of *lecF*, *lecX*, and cell density control genes, *epsB* and *iolG*. Gene expression in the root interior and stem is shown on a base-2 logarithmic scale relative to expression in GMI1000 cells on the rhizoplane. Asterisks indicate a difference in gene expression between the stated condition and the rhizoplane (Student’s t-test; *P≤0.05, **P≤0.01). **B** and **C**) **LexF and LecX negatively modulate *Rps* attachment to the root surface.** Roots of 4-day-old seedlings were inoculated with 10^4^ CFU of wild-type (gray), Δ*lecF* (light blue), Δ*lecF+lecF* (complemented mutant, striped blue), Δ*lecX* (light green), or Δ*lecX+lecX* (complemented mutant, striped green). At 2 hpi roots were washed, homogenized, and dilution plated to quantify the rhizoplane population. Each symbol represents four pooled roots. Data shown reflect three experiments, each with 6-10 technical replicates per treatment. Asterisks indicate a difference between wild-type and the lectin mutants and complemented strains (B, Krusal-Wallis test, Δ*lecF*: *P*=0.0089, Δ*lecX*: 0.0089; C, Mann-Whitney, *P*=0.0008). **D, E,** and **F) Mutating both *lecF* and *lecX* increased *Rps* root attachment and virulence but did not affect colonization of tomato roots or stems.** Roots of 4-day-old tomato seedlings were inoculated with 10^4^ CFU of *Rps* GMI1000 wild-type or the Δ*lecF/*X double mutant. After 48 h, roots were surface sterilized, homogenized, and dilution plated. Experiments were repeated three times with 9 to 12 technical replicates per treatment (Mann-Whitney test, *P*=0.60). Horizontal yellow bars indicate the geometric mean. **E)** 21-day-old tomato plants were inoculated through a cut petiole with 2000 CFU of wild-type or Δ*lecF/*X *Rps*. At 3 dpi, mid-stem samples above the point of inoculation were harvested, homogenized, and dilution plated. Data shown are from three experiments, each with 11-15 plants per treatment (Mann-Whitney test, *P*=0.95). The numbers above the x-axis denote the total number of uncolonized plants. **F)** 21-day-old tomato plants were soil soak inoculated with 50 mL of 10^8^ CFU/mL wild-type GMI1000 or Δ*lecF/X*. Disease severity was rated over 14 days on a scale from 0 (no wilting) to 4 (76-100% of plant wilting). Each point indicates the area under the disease progress curve for one plant. Data shown are from three experiments, each with 11-15 plants per treatment (Student’s t-test, *P*=0.0201).

*Rps* strongly upregulated *lecF* and *lecX* in both the root endosphere and stem, with at least a 100- fold higher expression than in bacteria on root surfaces (Fig 1A).

To determine if LecF and LecX help *Rps* colonize the root endosphere, we inoculated tomato seedling roots with 10,000 CFU of either wild-type strain GMI1000, *ΔlecF*, *ΔlecX*, or *ΔlecF/X*, a double mutant lacking both *lecF* and *lecX*. We also assessed contributions of LecF and LecX to stem colonization by introducing 2000 CFU of wild-type or lectin mutant strains directly into the vascular tissue through a cut leaf petiole, bypassing the root phase of disease. There were no differences in population sizes between wild-type *Rps* and the lectin mutants colonizing tomato root endospheres or mid-stems (Fig 1D and E, Fig S1A-D). The single lectin mutants were also as virulent on whole plants as the wild-type (Fig S1E and F). While *ΔlecF/X* and the wild-type both reached a final disease index near 4 (all plants dead), the lectin double mutant caused disease at a slightly but significantly faster rate (Fig 1F and S1G).

Although lectin gene expression was relatively low in *Rps* cells on the rhizoplane, we tested the hypothesis that *lecF* and *lecX* together help the bacterium attach to roots.

Unexpectedly, *ΔlecF*, *ΔlecX*, and *ΔlecF/X* all hyper-attached to roots, with the loss of *lecX* having the greatest effect (Fig 1B and C). Complementation of the single lectin mutants, Δ*lecF+lecF* and Δ*lecX+lecX*, restored near wild-type levels of adhesion to roots. Taken together, these plant assays indicate that while LecF and LecX are not required for wild-type tomato colonization or virulence, the lectins do constrain *Rps* attachment to tomato root surfaces.

### Lectin mutants hyper-attach in static conditions but are deficient in attachment under flow

Because lectins modulated bacterial attachment to tomato roots under static conditions, we further evaluated the role of *lecF* and *lecX* in static biofilm formation *in vitro*. The classic polyvinyl chloride (PVC) plate assay indirectly quantifies bacterial biofilm adhered to well walls via crystal violet staining. We also used confocal microscopy to visualize bacterial aggregates formed at the bottom of a cover glass chamber where the oxygen concentration is low. This method reveals bacterial aggregate structure and cohesion behaviors.

Intriguingly, all three lectin mutants formed three-fold more biofilm than the wild-type on PVC plates (Fig 2A). The magnitude of biofilm increases did not differ among the mutants, suggesting that LecF and LecX may be functionally redundant. Complementing the single lectin mutants with a single copy of the deleted gene restored wild-type levels of biofilm formation on PVC plates (Fig 2A). These results indicate that individually and together LecF and LecX negatively modulate *Rps* biofilm formation in stasis. On cover glass, wild-type GMI1000 formed irregular aggregates across the surface. Consistent with the PVC plate experiments, the lectin mutants formed fewer but much larger and thicker aggregates than wild-type GMI1000 (Fig 2B- E). These *in vitro* biofilm assay results were consistent with the hyper-attachment of *ΔlecF*, Δ*lecX,* and Δ*lecF/X* to tomato root surfaces.

**Figure 2.**
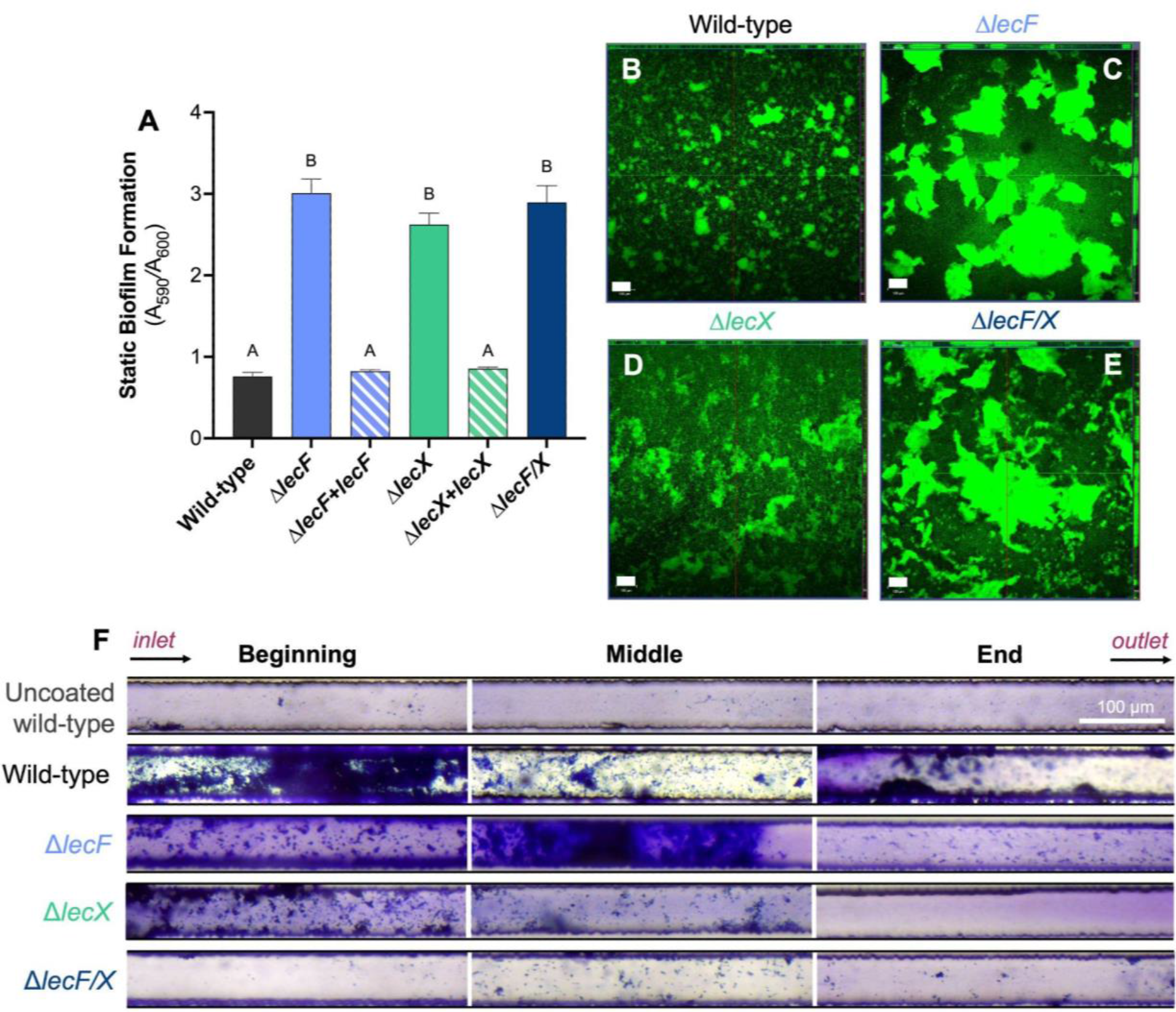
Effects of deleting *Rps* lectins on biofilm formation *in vitro*. A and B) LexF and LecX constrain *Rps* biofilm formation in static conditions. A) 10^7^ CFU/mL of wild-type GMI1000, Δ*lecF,* Δ*lecF+lecF (*complemented mutant), Δ*lecX,* Δ*lecX+lecX (*complemented mutant), and Δ*lecF/*X in CPG broth were aliquoted into a 96-well PVC plate. After 24 h, biofilms were stained with 1% w/v crystal violet and measured at A_590nm_. Data are from three independent experiments, each with 16-32 technical replicates per treatment. Different letters over bars indicate differences among strains as determined by ANOVA (*P*<0.0001). **B-E)** 10^7^ CFU/mL wild-type GMI1000, Δ*lecF,* Δ*lecX,* or Δ*lecF/*X suspended in CPG broth were aliquoted into 8-well chambered coverglass slides. Cultures were grown statically for 3 days at 28°C with fresh media added daily and cells were stained with SYTO9 before confocal imaging. Representative images show the orthogonal view at the middle of biofilm Z-stacks. Experiments were repeated twice with 3-4 technical replicates per treatment. The white scale bar indicates 100 µm. **F) LecF and LecX are essential for biofilm formation in xylem sap under flow.** 50x50 µm-cross-section microfluidic channels were coated with carboxymethyl cellulose-dopamine (CDC-DOPA). *Rps* wild-type GMI1000, Δ*lecF,* Δ*lecX,* and Δ*lecF/X* were suspended at 10^9^ CFU/mL in *ex vivo* tomato xylem sap, seeded into channels for 6 h and then incubated for 3 days at a flow rate of 38 µL/h. Biofilms were stained with 1% crystal violet and imaged with a light microscope. The experiment was repeated twice and representative images are shown. White bar indicates 100 µm.

A possible explanation for the sticky behavior of the *lecF* and *lecX* mutants is that *Rps* compensates for the loss of *lecF* or *lecX* by upregulating the remaining lectin genes, especially *lecM*. To test this hypothesis, we measured expression of all three lectin genes in the lectin mutant backgrounds. In Δ*lecF*, *lecX* gene expression trended higher but was not significantly different from wild-type, while *lecF* was upregulated in Δ*lecX.* In both single mutants, *lecM* expression was slightly elevated, but it was unchanged in the Δ*lecF/X* double mutant (Fig S2). Thus, upregulation of *lecM* only in the single mutants cannot explain the increased attachment behaviors of Δ*lecF/X*.

In addition to the static biofilm experiments discussed above, we also evaluated biofilm formation in a dynamic flowing environment using a recently-developed microfluidic system that mimics tomato xylem [48]. Channels were coated with the primary plant cell wall component cellulose using carboxymethyl cellulose-dopamine. Without this cellulose coating, the wild-type could not form mature biofilms in the microfluidic channels (Fig 2F). Bacteria were grown in this system under the flow of *ex vivo* tomato xylem sap. The lectin mutants behaved very differently in this flowing system than they did in the static seedling root, PVC plate, and cover glass systems. In cellulose-coated channels the wild-type strain formed thick biofilms that spanned the diameter of the channel at the beginning, with thinner biofilms observed toward the end. In contrast, the *lecF* and *lecX* mutants formed mostly small, diffuse aggregates, with the loss of *lecX* appearing to have a greater effect. While Δ*lecF* aggregation was observed at the middle and end of the channel, Δ*lecX* attachment was absent near the outlet.

Under shear stress, the loss of both *lecF* and *lecX* had a striking additive effect that nearly abolished biofilm development. Only a few aggregates were visible throughout the channel, indicating that without LecF and LecX, *Rps* cells failed to adhere, cohere, and develop biofilms in a xylem-mimicking environment.

### Heterologous expression of lectins in non-native strains suppresses biofilm formation *in vitro*

About 60% of genes in RSSC strain genomes belong to the core genome; this includes *lecM* [49]. Bioinformatic analysis of 393 sequenced strains in the RSSC revealed that while all strains have *lecM*, only ∼20% of phylotype IIB strains have *lecF,* ∼60% of phylotype IIA strains have *lecF*, and no sequenced strains in phylotype III have *lecX* (Fig S1). Only seven RSSC strains have *lecM* alone, suggesting that *lecF* and/or *lecX* confer additional, possibly different, benefits from *lecM*. Interestingly, three non-plant pathogenic *Ralstonia* spp.—*R. mojiangensis*, *R. mannitolilytica*, and *R. pickettii—*encode versions of LecF that are 91% identical at the amino acid level with LecF of the *Rps* model strain GMI1000, the primary strain used in this paper (Fig 3A).

**Figure 3.**
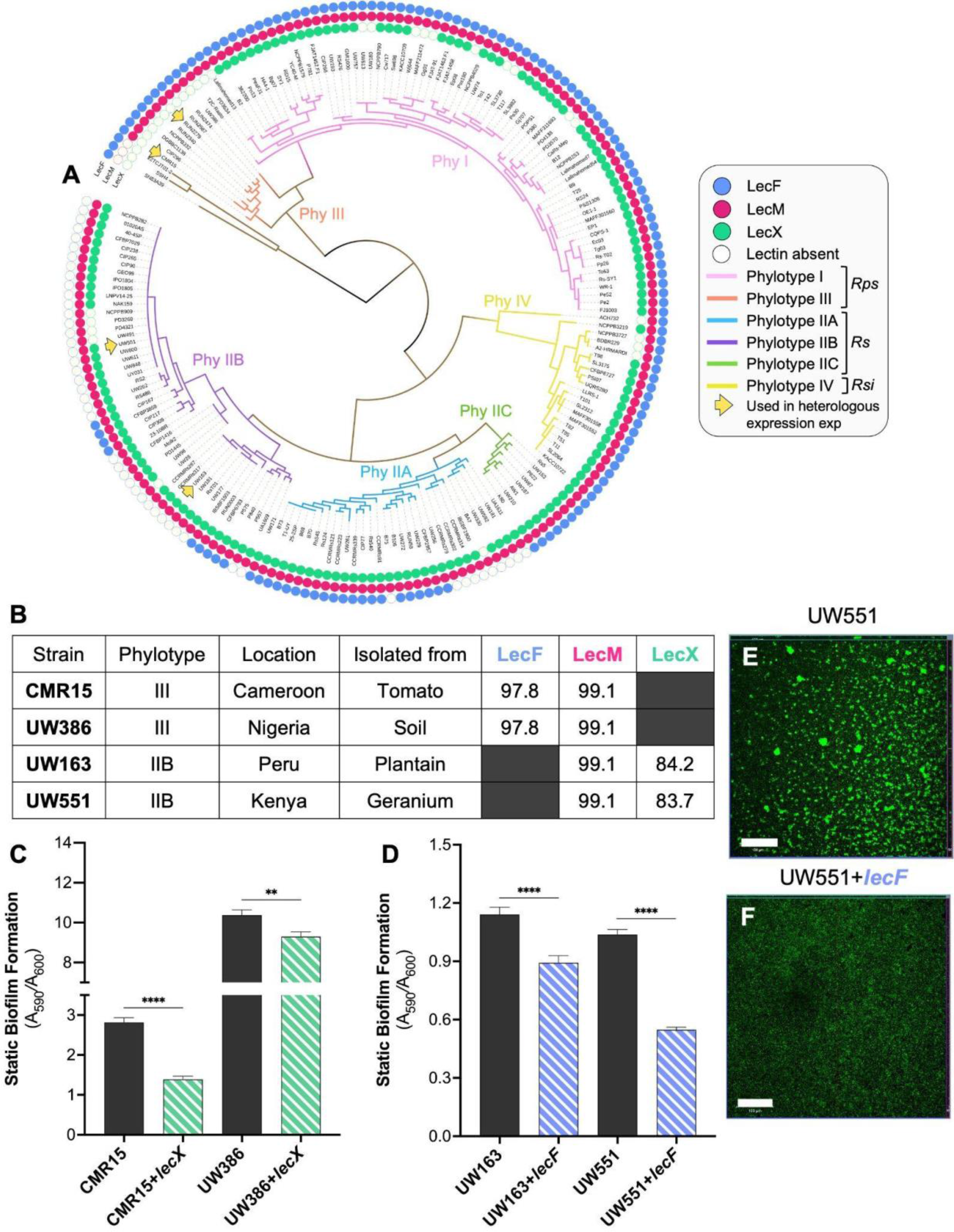
Heterologous expression of GMI1000 *lecF* and *lecX* in other plant pathogenic *Ralstonia*. **A)** Phylogenetic tree of the RSSC was constructed using 49 conserved genes and the RSSC Phylogenomics narrative on KBase [76,77]. Complete genomes of 187 RSSC strains and 3 other *Ralstonia* spp. were included in this analysis. Protein BLAST was used to determine lectin conservation and the outermost circles indicate the presence (filled icon) or absence (open icon) of *lecF* (blue), *lecM* (red) or *lecX* (green). Strains marked with yellow arrows were used for heterologous expression experiments. **B) GMI1000 *lecF* and *lecX* were expressed in four *Rs* and *Rps* strains that naturally lack them**. Strain characteristics and identity or similarity with GMI1000 lectin proteins are shown. Dark gray boxes indicate the gene is absent from that strain. **C-F) Heterologous expression of *lecF* and *lecX* in *Rs* Phylotype II and *Rps* Phylotype III strains reduced static biofilm formation. C** and **D)** *Ralstonia* strains CMR15, CMR15+*lecX*, UW386, UW386*+lecX,* UW163, UW163+*lecF*, UW551, and UW551*+lecF* were suspended at 10^7^ CFU/mL in CPG broth, incubated in 96-well PVC plates for 24 h, stained with crystal violet, and biofilm was quantified as A_590nm_. For UW386, data reflect four independent experiments, each with 12-48 technical replicates. For CMR15, UW163, and UW551 strains, data represent three independent experiments with 12-48 technical replicates. Asterisks indicate a difference between the wild-type and heterologous-expression strains (Student’s t-test; C, *P*<0.0001, *P=*0.0027; D, *P*<0.0001). **E** and **F)** 10^7^ CFU/mL wild-type UW551 and UW551+*lecF* suspended in CPG broth were aliquoted into 8-well chambered coverglass slides and cultured. statically for 3 days at 28°C and then stained with SYTO9 for confocal imaging. Representative images show the orthogonal view at the middle of biofilm Z-stacks. Experiments were repeated twice with 2-4 technical replicates per treatment. The white scale bar indicates 100 µm.

Although strain GMI1000 has all three lectin genes, many RSSC strains have *lecM* plus either *lecF* or *lecX*. To explore if GMI1000 LecF and LecX are functionally redundant and see if they can negatively impact static attachment in other *Ralstonia* strains, we constructed multiple *Rs* and *Rps* strains that heterologously express the absent lectin. *LecX*_GMI1000_ was added to two phylotype III strains: *Rps* UW386, a soil isolate from Nigeria, and *Rps* CMR15, a tomato isolate from Cameroon (Fig 3B). Similarly, *lecF*_GMI1000_ was expressed in two phylotype IIB strains: UW551, a geranium isolate from Kenya, and UW163, a plantain isolate from Peru (Fig 3B). qRT-PCR confirmed that the heterologous *lecX* or *lecF* were highly expressed in these constructed strains (Fig 4B and C). As described above, individually mutating *lecF* and *lecX* in GMI1000 dysregulates the expression of the remaining lectins (Fig S2). While *lecF*_UW386_ expression was slightly increased in UW386-*lecX*, heterologous expression of *lecX*_GMI1000_ in CMR15 and l*ecF*_GMI1000_ in UW163 and UW551 did not change the expression of the strains’ native lectins (Fig S4).

**Figure 4.**
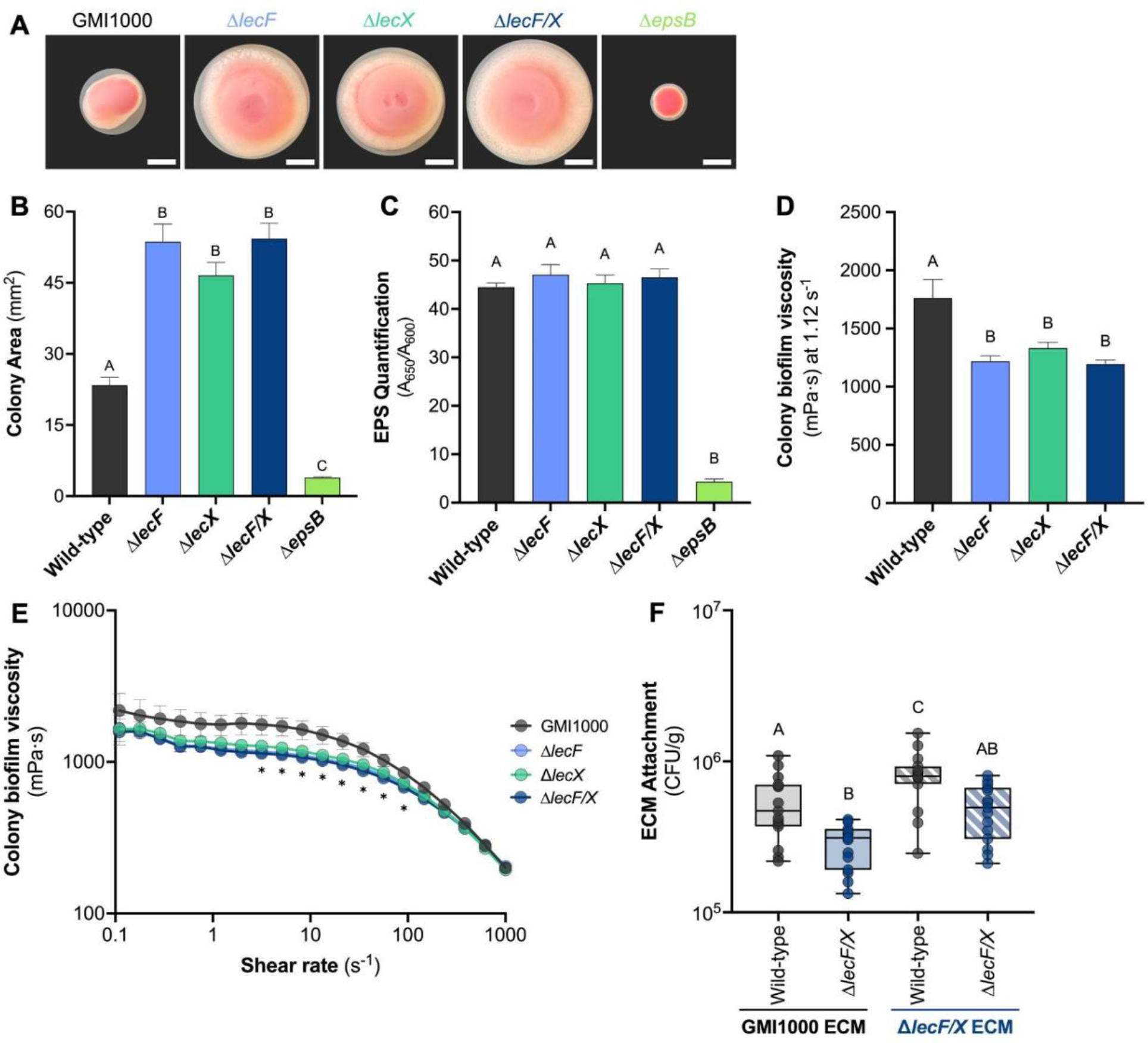
Effect of *Rps* lectins on the bacterial extracellular matrix. A-B) LecF and LecX are required for normal colony coherence. 30 CFU of wild-type *Rps* GMI1000, Δ*lecF,* Δ*lecX,* Δ*lecF/X,* or Δ*epsB* were spread onto CPG media supplemented with tetrazolium chloride. Plates were photographed after 4 days’ incubation at 28°C. Representative colony images are shown in A and the white scale bar indicates 2 mm. B) Colony area was measured using ImageJ. Data shown reflect two independent experiments containing 14-31 technical replicates. Different letters above bars indicate differences among the strains (ANOVA, *P*<0.0001). **C) *Rps* lectin mutants produce wild-type levels of EPS I.** 10^8^ CFU of wild-type GMI1000, Δ*lecF,* Δ*lecX*, Δ*lecF/X*, and Δ*epsB* were spread onto CPG plates. Following a 4-day incubation at 28°C, bacteria were scraped from plates and resuspended in water. 100 µL of this bacterial suspension was aliquoted into 96-well plates coated with Agdia anti-EPS-I monoclonal antibodies and DAS- ELISA was performed. Data shown reflect two independent experiments, each containing 6-8 technical replicates. Different letters above bars indicate differences among the strains (ANOVA, *P*<0.0001). **D** and **E**) **Lectins contribute to wild-type viscosity of the *Ralstonia* extracellular matrix (ECM)**. 1 mL of colony biomass was scraped from 2% w/v agar plates following a three day incubation at 28°C. The biofilm colony viscosity of wild-type GMI1000, Δ*lecF*, Δ*lecX*, Δ*lecF/X* was measured at strain rates ranging from 0.01 to 1000 s^-1^ and the mean viscosity at a shear rate of 1.12 s^-1^ is displayed in panel D (D,ANOVA, P=0.006; E, Repeated measures Two- way ANOVA, *P*<0.0001). **F) Crude ECM from a Δ*lecF/X* double mutant was more adhesive than wild-type ECM, and LecF and LecX contribute to *Rps* attachment to ECM**. Mixed cellulose ester (MCE) membranes were incubated in crude ECM extract from wild-type GMI1000 (solid box) or Δ*lecF/X* (striped box). Membranes were washed and incubated with a suspension of wild-type GMI1000 (gray) or Δ*lecF/X* (dark blue). After 1 h, membranes were gently washed, homogenized, and dilution plated to quantify the adhering bacteria. Data shown reflect three experiments with 5 technical replicates each. Different letters indicate significant differences (ANOVA, *P*<0.0001).

Because mutating *lecF* and *lecX* increased GMI1000 biofilm formation in static conditions (Fig 2A), we hypothesized that expressing non-native lectins in strains that naturally lack them would reduce biofilm formation. For all four strains evaluated, adding *lecF* or *lecX* did in fact suppress biofilm formation on PVC plates (Fig 3C and D). The greatest effect was in UW551+*lecF* and CMR15+*lecX*, where an additional heterologous lectin reduced biofilm formation about 50%. Visualizing biofilm architecture of heterologous-expression strains with confocal microscopy revealed that UW551+*lecF* cells failed to cohere and form aggregates. Wild-type UW551 forms spherical clusters on glass and these structures were notably absent in UW551+*lecF* (Fig 3E and F). These results are consistent with the biofilm behavior of GMI1000 Δ*lecF*, Δ*lecX*, and Δ*lecF/X* mutants, suggesting that in static conditions, LecF and LecX negatively modulate biofilm formation in plant pathogenic *Ralstonia* spp.

### Lectins influence the behavior and biomechanics of the ECM

On agar plates, colonies of all three lectin mutants, Δ*lecF*, Δ*lecX*, Δ*lecF/X*, appeared atypically loose and spread more than wild-type GMI1000, forming colonies that were more than twice the area of the wild-type after 4 days of growth (Fig 4A and B). The *epsB* mutant, which secretes very little exopolysaccharide, had a characteristically dry, small, and round colony morphology. This control mutant confirmed that self-produced polysaccharides are a significant proportion of the GMI1000 colony biomass and presumably the ECM in biofilms.

Mutation of *lecM*, the third lectin in GMI1000, altered the production and expression of the exopolysaccharide biosynthetic cluster in a closely related phylotype I strain [42,50]. To determine if EPS production was similarly altered in Δ*lecF* and Δ*lecX*, we quantified EPS in colonies with DAS-ELISA using antibodies specific to EPS-I from members of the RSSC. As expected, the *epsB* mutant control produced about 10-fold less EPS-I than the wild-type. The wild-type and lectin mutants produced the same amount of exopolysaccharide (Fig 4C). Thus, the spreading colony morphology and enhanced attachment behaviors of the lectin mutants cannot be explained by altered production of polysaccharides.

Additionally, the lectin mutant colonies were surrounded by a larger ring of white material, presumably EPS, at the colony edge and they formed craters at their centers that suggested reduced structural integrity. Gradients in extracellular polymer concentration could guide both osmotic spread of and viscous flow within biofilms [51,52]. Our observations thus suggest a link between the lectins and the mechanical properties of the biofilm.

Given the abnormal spreading morphology of Δ*lecF/X* colonies, we hypothesized that the ECM of lectin mutants is physically different from that of the wild-type. To test this, we measured the shear viscosity of the colony biofilm over increasing shear rates for Δ*lecF,* Δ*lecX*, and Δ*lecF/X* (Fig 4E). While all the strains thinned under shear, the viscosity of the lectin mutants was consistently lower than that of the wild-type (Fig 4D and E). EPS production has been shown to generate outward forces via osmotic pressure gradients that overcome viscous resistance to colony spreading [52]. Our measured viscosities are consistent with such a mechanism of spreading.

We also assessed the behavior of the ECM by quantifying attachment of bacterial cells to an ECM-coated membrane [25]. Interestingly, GMI1000 attached to crude ECM from Δ*lecF/X* more than to ECM from wild-type colonies (Fig 4F). The loss of *lecF* and *lecX* together increased the adhesiveness of the ECM, offering a possible explanation for the Δ*lecF/X* hyper-attachment behaviors in static conditions. Lectins have been shown to bind exopolysaccharides and stabilize the biofilm matrix encompassing other bacteria [2,24]. We hypothesized that *lecF* and *lecX* have a similar function in *Rps*. The lectin double mutant was deficient in attachment to the ECM-coated membranes, regardless of the ECM source strain (Fig 4F). Together, these data suggest that LecF and LecX contribute to both the ECM mechanical properties and bacterial interactions with the ECM.

### EPS I production is required for attachment under flow, but not under static conditions

EPS I, the major extracellular polysaccharide in the RSSC, is a virulence factor in both *Rs* and *Rps*; it is critical for colonization of the stem and vertical spread in the stem [33,35,36]. It has been assumed EPS I has a major role in *Ralstonia* biofilm development as it does in several well-studied bacteria and because it is a component of the biofilm ECM, but studies of static biofilm formation yielded mixed results [53,54]. To investigate this further in both static and flowing environments, we used an EPS I-deficient *epsB* mutant.

In *Rs* UW551, EPS contributes to tomato root attachment [11]. We hypothesized that *Rps* GMI1000 also requires EPS for host adhesion. However, in GMI1000 similar numbers of Δ*epsB and* wild-type cells adhered to the tomato rhizoplane (Fig 5A). Additionally, GMI1000 Δ*epsB* formed similar amounts of biofilm as the wild-type on PVC plates and at the bottom of glass wells (Fig 5B-D). This suggests that EPS plays different roles in *Rs* and *Rps*.

**Figure 5.**
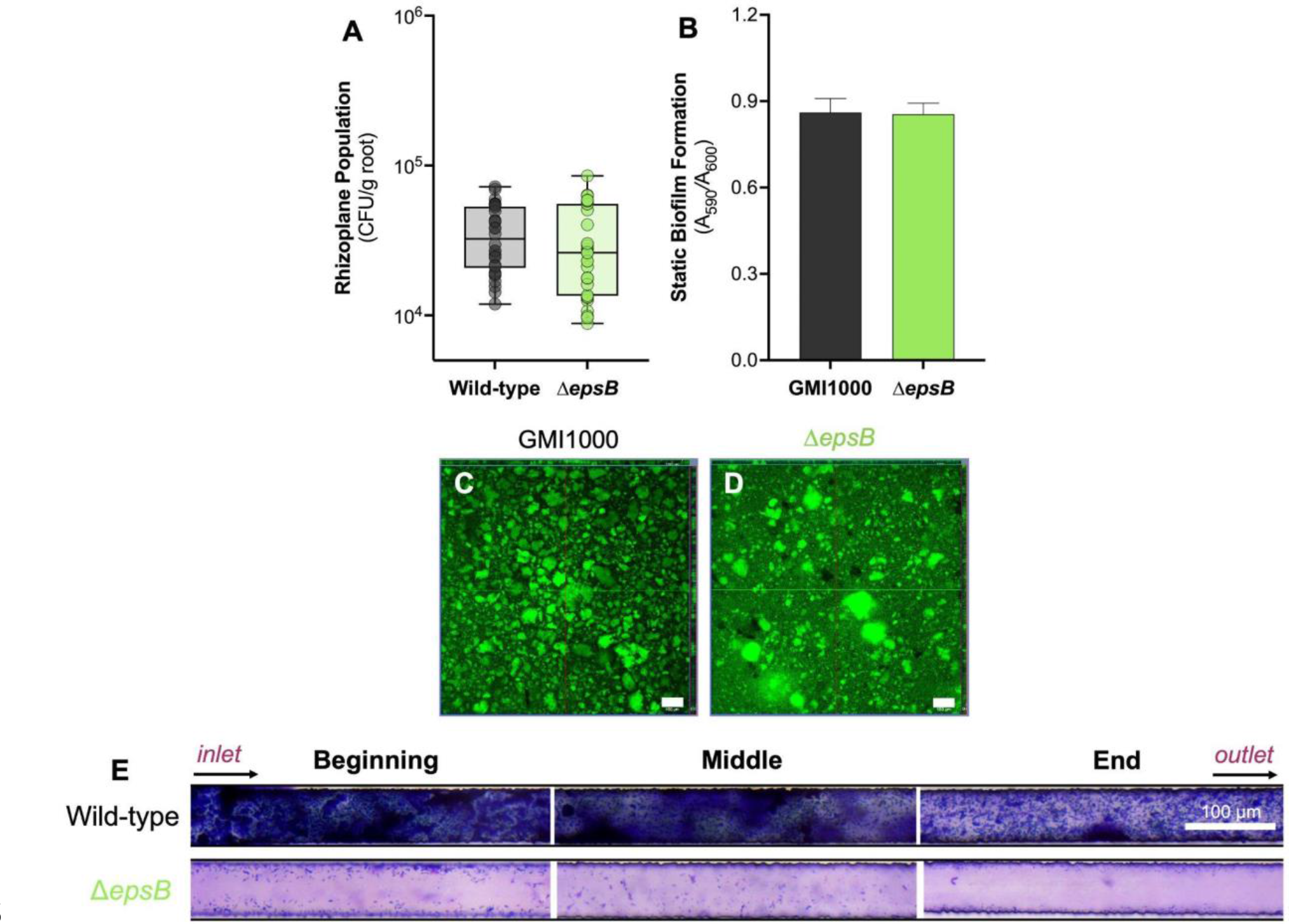
Effect of *Rps* EPS I on bacterial attachment. A-D) EPS I is not required for attachment to tomato roots, PVC, or glass under static conditions. A) Roots of 4-day-old seedlings were inoculated with 10^4^ CFU of wild-type or Δ*epsB* for 2 h, then washed, homogenized, and dilution plated to quantify the rhizoplane population. Each dot represents four pooled roots. Data are shown from three experiments each containing 10 technical replicates (Mann-Whitney test, *P*=0.2979). **B)** 10^7^ CFU/mL GMI1000 and Δ*epsB* in CPG broth were incubated in a 96-well PVC plate for 24 h without shaking. Biofilms were stained with 1% crystal violet and measured at A_590nm_. Data shown reflects five independent experiments with 16- 32 technical replicates (Student’s t-test; *P*=0.92). **C** and **D)** 10^7^ CFU/mL suspensions of wild-type GMI1000 (C) and Δ*epsB* (D) were incubated without shaking in 8-well chambered coverglass slides for 3 days at 28°C and then biofilms were stained with SYTO9 for confocal imaging. Representative images show the orthogonal view at the middle of biofilm Z-stacks. Experiments were repeated twice with 3-4 technical replicates each. The white scale bar indicates 100 µm. **E) EPS I is essential for *Rps* biofilm formation under flow.** 50x50 µm- cross-section microfluidic channels coated with carboxymethyl cellulose-dopamine (CDC- DOPA) were seeded for 6 h with *Rps* wild-type GMI1000 or Δ*epsB* suspended at 10^9^ CFU/mL in *ex vivo* Bonny Best xylem sap. Xylem sap was then pumped through devices for 3 days at a flow rate of 38 µL/h. Biofilms were stained with 1% crystal violet and imaged with a light microscope. The experiment was repeated twice and representative images are shown. The white bar indicates 100 µm.

While *Rps* EPS I was not critical for static biofilm formation or root attachment, it might nevertheless function in attachment under shear stress. We tested this hypothesis using the cellulose coated microfluidic channels containing tomato xylem sap, as described above. Under flow, Δ*epsB* failed to form mature biofilms; similar to Δ*lecF/X*, the EPS I-deficient mutant made only small and diffuse aggregates over the length of the microfluidic channel (Fig 5E). Together these results show that EPS I contributes to *Rps* adhesion and biofilm development specifically in flowing conditions.

## Discussion

We explored the functions in the RSSC of two highly expressed lectins, LecF and LecX, and the major exopolysaccharide EPS I in static and flowing environments. GMI1000 has a third lectin, LecM, a known virulence factor in *Rps* and *Rs* and contributor to static biofilm formation *in vitro* [41,42]. The presence of *lecM* in all the ∼400 RSSC strains analyzed indicates it has a core function in the life cycle of plant pathogenic *Ralstonia*, regardless of the plant host or method of transmission.

Unlike for *lecM*, mutating *Rps lecF* and *lecX* enhanced bacterial attachment in static environments, both on the plant root and *in vitro* [41,42]. Consistent with this, heterologous expression of *lecX* or *lecF* in RSSC strains that naturally lack them reduced biofilm formation. Together these results suggest that LecF and/or LecX prevent wild-type cells from excessive attachment to plant surfaces or to each other in the absence of flow, thereby promoting dispersal. We demonstrate that wild-type *Rps* specifically represses *lecF* and *lecX* gene expression at the rhizoplane, possibly to enhance root attachment. Additionally, the reduced expression of *lecF* and *lecX* in the Δ*phcA* may explain the sticky phenotype of this quorum sensing mutant on host roots [29]. LecM does not function in this initial host interaction [41]. Our results demonstrate LecF and LecX have a distinct function from *L*ecM in static root attachment and biofilm formation.

The hyper-attachment behaviors of the lectin mutants were surprising, so we wondered if other indirect effects of mutating lectins could explain the excessive attachment of Δ*lecF,* Δ*lecX,* and Δ*lecF/X*. We tested two hypotheses: H_0_1, the mutation of *lecF* and *lecX* increased expression of *lecM*, resulting in hyper-attachment and H_0_2, the loss of *lecF* and *lecX* increased production of extracellular polysaccharides causing atypically loose colony morphology and enhanced attachment. Expression of *lecM* was unchanged in Δ*lecF/X* and all three lectin mutants produced wild-type levels of EPS, disproving these hypotheses.

Although the lectin mutants attached more in static conditions, the loss of *lecF* and *lecX* had the opposite effect under flow conditions that resemble the xylem habitat. LecF and LecX were required for biofilm formation in cellulose-coated microfluidic channels. These lectins may mediate *Rps* adhesion to cellulose, a component of xylem vessel walls, as well as *Rps* aggregation [43]. Our data suggest that LecF and LecX are functionally redundant under static conditions, but these two lectins contributed additively to biofilm development under flow.

While LecM may enable biofilm formation in multiple environments, we discovered a flow-specific role in attachment for LecF and LecX.

Lectin mutants had opposing attachment phenotypes in static conditions (rhizoplane, glass slide, and PVC plate) and under flow. These biochemically and physically distinct environments elicited very different bacterial attachment behaviors. Some bacteria and lectins display force- and flow-enhanced attachment (so-called catch bonds) to ligands [55]. One of the most well-studied models is FimH, a mannose-binding lectin located at the tip of the type I pilus in *Escherichia coli* [56,57]. As shear stress increases, FimH mediates 100-fold greater *E. coli* adhesion. Under static or very low flow conditions, FimH rapidly disassociates from its ligand, enabling bacterial motility and dispersal [56,57]. If LecF and LecX are catch-bond-forming adhesins, it may explain their evidently force-dependent role in attachment. Additional flow cell experiments should characterize *Rps* responses to hydrodynamic forces and the intersecting influence of lectin-polysaccharide interactions.

LecF is present in some non-plant pathogenic *Ralstonia* spp., but LecX and LecM are not. We speculate LecF mediates eukaryotic and*/*or interbacterial interactions that are not unique to the plant host or environment. LecF forms a six-bladed β-propeller architecture similar in structure and sequence to the fucose-binding lectin BambL in *Burkholderia ambifaria* [58–60]. LecF and BambL, which is 76% identical to the *Rps* lectin at the amino acid level, both have three intramonomeric and three intermonomeric glycan binding sites and very similar specificities for fucosylated oligosaccharides [37,58]. *In silico* mutagenesis revealed that residues Arg17, Glu28, Trp76, Trp81 in intramonomeric binding sites are crucial for LecF interactions with Me-α-L-fucoside [61]. In the opportunistic human pathogen *B. ambifaria*, BambL functions as a B cell superantigen binding fucose on blood B cell receptors, possibly triggering activation-induced cell death and disrupting the adaptive immune response [62,63]. Lectin interactions with surface glycans on eukaryotic hosts are evidently important for pathogenesis. Lectin binding to bacterial polysaccharides can also mediate cell-cell interactions driving biofilm development, like with RSSC LecM.

Lectins and secreted enzymes in the ECM influence the stability, structure, and physiology of biofilms [24,28,64]. We hypothesized that LecF and LecX alter the properties of the ECM, contributing to the opposing biofilm phenotypes in static and flow conditions.

Extracted ECM from Δ*lecF/X* was more adhesive than that of the wild-type in static conditions, which may explain the lectin mutant hyper-attachment. Importantly, the lectins also contribute to ECM attachment; without them, the ECM may lose structural integrity, resulting in the spreading colony morphology of the lectin mutants.

While this loose matrix may yield more biofilm at stasis, it could be deleterious for biofilm development under mechanical stress as we observed with the lectin mutants in the artificial xylem system. Both LecF and LecX increase the viscosity of the colony biofilm, possibly supporting microcolony development and expansion in a flowing environment [65]. ECM polymers have been shown to determine the viscoelastic properties of biofilms [65,66]. Psl, one of three major exopolysaccharides in *P. aeruginosa*, enhanced cross-linking and elasticity promoting a stiffer biofilm matrix [67]. Here we show a proteinaceous component of biofilms also influences the biomechanics of the ECM. Analysis of the cellular localization of LecF and LecX, direct protein-EPS interactions, and spatial heterogeneity in extracellular polymer concentration is needed to further elucidate the role of lectins in the biofilm matrix.

EPS is critical to biofilm development for many bacteria, and it is a key component of the biofilm matrix [1,2,68]. While RSSC strains produce copious amounts of EPS *in vitro* and in the stem at high cell densities, EPS has not been directly visualized in biofilms *in planta* [33,69]. A recent study on endoparasitism of *Fusarium oxysporum* showed EPS I is a core component of *Rs* biofilms on fungal hyphae [54]. In *Rps* OE1-1, EPS I contributes to biofilm formation on PVC plates but contrarily, an EPS I-deficient mutant formed a similar number of mature biofilms in *ex vivo* apoplastic fluid [53,54]. We provide evidence that *Rps* GMI1000 EPS I is particularly essential for biofilm development in flowing conditions, as are LecF and LecX.

During colonization, plant pathogenic microbes attach to the host and form biofilms in diverse microenvironments. The foliar pathogen *Xanthomonas fuscans* subsp. *fuscans* forms biofilms on bean seeds, leaf surfaces, and in the leaf apoplast, while the insect-vectored pathogen *X. fastidiosa* forms biofilms in the precibarial canal of sharpshooter leafhoppers and the plant xylem [70,71]. Nearly all RSSC bacteria begin their life cycle at the rhizosphere, aggregating on and in the root and ultimately, the xylem.

We propose that *Rps* lectins LecF and LecX function very differently depending on the physical environment (Fig 6). When in relatively static environments like the rhizosphere or an occluded xylem vessel, LecF and LecX constrain bacterial attachment to enable the pathogen to spread to new locations. While in the flowing environment of healthy xylem vessel, LecF and LecX interactions with plant cell walls and ECM facilitate bacterial adhesion and biofilm maturation under shear stress. Biophysical studies could determine if LecF and LecX respond to mechanical cues to help *Rps* cells balance dispersal and attachment over the complex bacterial wilt disease cycle.

**Figure 6.**
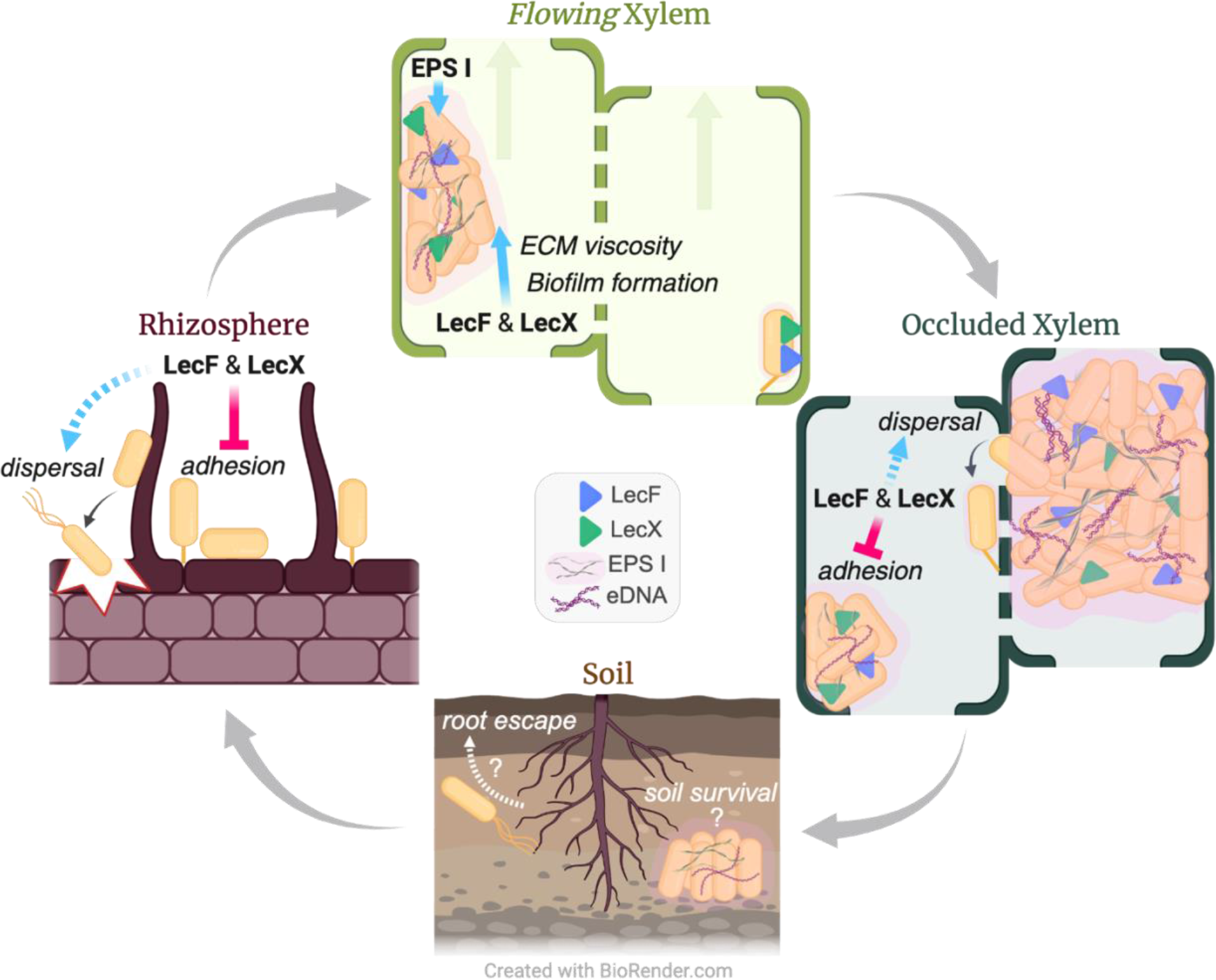
Proposed roles of EPS I, LecF, and LecX in bacterial wilt disease. Data presented here suggest that in the relatively static environment of the host rhizoplane, the lectins constrain *Rps* adhesion, thereby promoting pathogen dispersal in the rhizosphere and root invasion. Once inside the plant, *Rps* colonizes the water transporting xylem tissue. When *Rps* cells enter a healthy, flowing xylem vessel, LecF, LecX and EPS I mediate bacterial attachment to the vessel wall and to other bacteria. The lectins enable biofilm development by binding ECM polysaccharides. Under shear forces created by transpirational flow, the lectins increase the biofilm viscosity, supporting biofilm development and expansion. As the pathogen occludes vascular flow, creating a static environment, LecF and LecX once again negatively modulate attachment to facilitate *Ralstonia* dispersal to new vessels or escape from the host. The carbohydrate-binding lectins LecF and LecX thus have a novel environment-specific role in *Rps* attachment behaviors.

## Materials and methods

### Bacterial strains and mutants

*Escherichia coli* was grown at 37°C in LB and RSSC strains were cultured in CPG rich medium at 28°C with antibiotics when appropriate (15 mg/L tetracycline, 25 mg/L kanamycin, 25 mg/L gentamicin, 100 mg/L ampicillin). The bacterial strains, plasmids, and primers used in these experiments are listed in Table S1. Complete, unmarked deletions of *lecF* and *lecX* were made in *Rps* strain GMI1000 using the *sacB* positive selection vector pUFR80; lectin mutants were complemented using the chromosome insertion vector pRCK as previously described [72,73]. Briefly, mutagenesis vectors containing genomic regions up- and down-stream of *lecF* (pUFR80Δl*ecF*), *lecX* (pUFR80Δ*lecX*), *epsB* (pUFR80Δ*epsB*) were constructed using Gibson assembly. GMI1000 was transformed using natural transformation or electroporation and mutants were confirmed with PCR and sequencing [74]. The *lecF* mutant was transformed with pUFR80Δ*lecX* to generate the *ΔlecF/X* lectin double mutant. Gibson assembly was used to construct complementation and heterologous expression vectors containing the *lecF* (pRCK- *lecF*, pUC18-miniTn7T-*lecF*) or *lecX* (pRCK-*lecX,* pUC18-miniTn7T-*lecX*) ORF with the native promoter. To create heterologous-expression strains, *Rs* UW551 and *Rs* UW163 were transformed with pUC18-miniTn7T-*lecF* via electroporation [75]. *Rps* CMR15 and *Rps* UW386 were transformed with pUC18-miniTn7T-*lecX* via electroporation. Gene expression of *lecF* and *lecX* in heterologous expression strains was confirmed with qRT-PCR.

### Lectin conservation analysis

Using a modified version of the RSSC Phylogenomics narrative on KBase, 393 RSSC strains and three non-plant-pathogenic *Ralstonia* spp were assessed for LecF, LecM, and LecX protein conservation [76,77]. The threshold was defined as ≥75% amino acid identity over at least 90% of the protein sequence. Lectin protein sequences from *Rps* GMI1000 were used as the subject query in the KBase protein blast app to determine the calculated percent of strains containing LecF, LecM, and LecX. Of the analyzed RSSC strains, 187 were used to construct a phylogenetic tree based on 49 conserved genes. The resulting Newick file was uploaded to iTOL and edited to denote phylotypes (colored branches) and lectin conservation in each strain (colored open or filled circles) [78].

### *In vitro* and *in planta* gene expression analyses

Using the previously described hot phenol-chloroform extraction method, total RNA was collected from bacteria colonizing the rhizoplane, root endosphere, and mid-stems of Bonny Best tomato plants [10,40]. For the root phase of colonization, five-day-old axenic seedlings were flood-inoculated with 15 mL (rhizoplane) or 3 mL (root endosphere) of wild-type *Rps* GMI1000 at a low cell density (5x10^6^ CFU/mL) in chemotaxis buffer (10 mM potassium phosphate, pH 7.0, 0.1 mM EDTA, 1 mM MgSO4) [8]. Root rhizoplane bacterial populations were harvested 6 hpi and gently washed once in water to remove unattached cells. Seedling roots were cut from the hypocotyl and 40-50 roots were pooled into one sample for RNA extraction. This was repeated three times to obtain three independent replicates. To obtain root endosphere populations, roots were collected 48 hpi, surface sterilized with 30 s treatments of 10% bleach and 70% ethanol, and 6 roots were pooled per biological replicate. The experiment was repeated twice. To measure bacterial populations in mid-stems, 21-day-old plants were petiole-inoculated with 2 µL of 10^6^ CFU/mL GMI1000 and 3 dpi stem slices were collected above the site of inoculation. Four stem samples were pooled per biological replicate and the experiment was repeated three times. RNA was reverse transcribed into cDNA with the SuperScript™ VILO™ cDNA Synthesis Kit (Invitrogen). A targeted qRT-PCR measured expression of *lecF, lecM, lecX, iolG*, and *epsB* and two normalization genes (*rplM* and *serC*) using primers in Table S1. The ΔΔCt method was used to calculate relative gene expression during root endosphere and stem colonization in comparison to the rhizoplane. Log2 fold change values were graphed and statistical significance was determined with Student’s t-tests.

To compare lectin gene expression in the lectin mutants to the wild-type, cultures were grown in rich media to an OD_600nm_ between 0.2 and 0.6 (log phase). Cells were pelleted in stop solution (5% phenol in ethanol) and total RNA was extracted using the hot phenol-chloroform method and an analysis of lectin gene expression was conducted as described above.

### Plant virulence and stem colonization assays

Plant assays were performed as described [79]. Briefly, to evaluate disease severity in tomato, unwounded 21-day-old wilt-susceptible tomato cv. Bonny Best plants were soil soak inoculated with 50 mL of a 10^8^ CFU/mL water suspension of wild-type strain GMI1000, Δ*lecF*, Δ*lecX*, and Δ*lecF/X* and symptom development was rated on a scale from 0 (no wilting) to 4 (76- 100% wilting) for 14 days. To quantify bacterial populations in the stem, 21-day-old plants were petiole-inoculated with 2 µL of a 10^6^ CFU/mL water suspension of wild-type, *ΔlecF, ΔlecX*, and *ΔlecF/X*. 100 mg stem sections were harvested 1 cm above the inoculated petiole, homogenized in water with a PowerLyzer bead beater, and serially dilution plated on CPG media supplemented with tetrazolium chloride to quantify CFU/g stem.

### Seedling root attachment and colonization

Bacterial attachment to axenic tomato seedling roots was measured as described [10].

Briefly, surface sterilized Bonny Best seeds were germinated and grown for 4 days on 1% water agar plates covered with a sterile paper filter. For root attachment assays, seedling roots were individually inoculated with 10 µL of 10^6^ CFU/mL (low cell density) or 10 µL of 2x10^7^ CFU/mL (high cell density) and incubated under white light for two hours. Roots were washed with sterile water for 10 s, cut from the hypocotyl, blotted dry, weighed, and homogenized in water with a PowerLyzer bead beater. Four roots were pooled per technical replicate. Homogenized root samples were serially dilution plated on CPG media supplemented with tetrazolium chloride to quantify CFU/g root.

To capture bacterial populations in the root endosphere, seedling roots were individually inoculated with 10 µL of 10^6^ CFU/mL and incubated at root temperature in a 12-hour photoperiod for two days. Roots were surface sterilized in successive treatments of 10% bleach for 30 s, 70% ethanol for 30 s, and four one-minute washes with sterile water. Roots were cut from the hypocotyl, blotted dry, homogenized in water with a PowerLyzer, and serially dilution plated on CPG media supplemented with tetrazolium chloride to determine the CFU/g root.

### Colony morphology analysis

30 µL of a 10^3^ CFU/mL water suspension of wild-type GMI1000, Δ*lecF*, Δ*lecX*, Δ*lecF/X*, or Δ*epsB* were spread onto CPG media supplemented with tetrazolium chloride, an indicator of respiration that creates the red color in *Ralstonia* colonies. After a three-day incubation at 28°C, plates were photographed and ImageJ was used to quantify the colony area (mm^2^). For magnified images of the colonies, plates were imaged with a dissecting light microscope.

### Extracellular polysaccharide quantification and ECM attachment

EPS production was quantified using the ELISA-based PathoScreen® Kit for *Ralstonia solanacearum* (Agdia Inc). To prepare samples for the kit, 100 µL of 10^9^ CFU/mL wild-type GMI1000, *ΔlecF, ΔlecX*, *ΔlecF/X,* and Δ*epsB* were spread onto CPG agar. Following a four-day incubation at 28°C, lawns of bacteria were scraped from the plate, resuspended in sterile water, and standardized to an OD_600nm_ of 0.01 (or 10^7^ CFU/mL). 100 µL of the bacterial suspensions were aliquoted into the provided microtiter plate and a DAS-ELISA with anti-EPS antibodies was performed as per the manufacturer’s instructions. The experiment was repeated twice.

Attachment to ECM-coated mixed cellulose esters (MCE) membrane (MF-Millipore, #VSWP02500) was evaluated using a modified protocol from Pradhan *et al.* [25]. 100 µL of a 10^9^ CFU/mL suspension of wild-type GMI1000 and *ΔlecF/X* were spread onto CPG media plates and incubated for four days at 28°C. Bacteria were resuspended in water, standardized to an OD_600nm_ of 15 (or 1.5x10^10^ CFU/mL), and centrifuged for 15 min twice at 8000 rpm. Cell-free ECM was precipitated from the supernatant overnight in four volumes of acetone with 20 mM NaCl, and redissolved in water. Sterile MCE membranes were incubated in 500 µL of crude EPS for one hour with shaking at 100 rpm. Membranes were washed for 15 min three times in phosphate-buffered saline with Tween (PBS-Tween) and allowed to air dry. The resulting EPS- coated MCE membranes were incubated in 500 µL of 10^9^ CFU/mL wild-type GMI1000 and *ΔlecF/X* for one hour. Following incubation with bacteria, membranes were washed once with sterile water, homogenized in water with a PowerLyzer bead beater, and serially dilution plated on CPG media supplemented with tetrazolium chloride to determine the CFU/g membrane.

### Shear rheology

A rheometer was used to measure the shear rheology of total colony biomass produced by wild-type GMI1000 and the lectin mutants. Each strain was cultured from freezer stocks on TZC plates for 48h at 28°C. A single colony was used to inoculate CPG broth, which was incubated overnight at 28°C with 250 rpm shaking. Culture densities were measured using a spectrophotometer, densities were adjusted with water to a final OD_600nm_ equivalent to 5x10^5^ CFU/mL, and 100 µL of this suspension was spread on four TZC plates with 2% w/v agar per strain and incubated at 28°C for 72 h. At least 1 mL of bacterial colony biomass was collected from each plate by wiping the agar surface with a glass rod. Samples were stored in closed microcentrifuge tubes and measured within 4 h of collection.

Shear viscosity of each sample was measured using an Anton Paar MCR 302 rotational rheometer with a 25 mm parallel plate (PP 25) geometry. About 1 mL of bacterial biomass was placed on the rheometer base plate. The PP 25 plate was lowered to a gap of 1 mm, excess sample was trimmed. Viscosity was measured at 25 strain rates, increasing logarithmically from 0.01 to 1000 s^-1^. To ensure the machine had reached steady state, each time point was measured for longer than the inverse of the corresponding shear rate.

### Biofilm assays

Static biofilm assays were performed as described [28]. Briefly, 150 µL of 10^7^ CFU/mL wild-type GMI1000, *ΔlecF, ΔlecX*, *ΔlecF/X, ΔlecF+lecF, ΔlecX+lecX,* GMI1000Δ*epsB*, UW551, or UW551Δ*epsB* resuspended in CPG were aliquoted into PVC microtiter plates. Plates were sealed with a Breathe-Easy sealing membrane (Diversified Biotech) and incubated at 28°C for 24 h. The cell density of each well was measured, and biofilms were stained with 1% crystal violet. Following three washes with sterile water, biofilms were de-stained with 200 µL of 95% ethanol and the resulting solution was measured at an OD_590nm_. The OD_590nmn_ data was standardized by cell density measured at OD_600nm_.

For static glass biofilm assays, 500 µL of 10^7^ CFU/mL GMI1000, *ΔlecF, ΔlecX*, *ΔlecF/X,* GMI1000Δ*epsB*, UW551, or UW551Δ*epsB* resuspended in CPG were aliquoted into the wells of an 8-well Nunc Lab-Tek II chamber coverglass (Thermo Fisher Scientific). The cover glass was covered with the provided cap and incubated at 28°C for 72 h. The spent medium was refreshed daily. Following incubation, biofilms were stained with SYTO9 per the instructions of the LIVE/DEAD Cell Imaging Kit (Invitrogen). Biofilms were imaged with a Zeiss LSM 710 laser scanning confocal microscope with appropriate filter sets for SYTO9 and propidium iodine at the University of Wisconsin-Madison Newcomb Imaging Center.

Biofilm formation under xylem-mimicking conditions was conducted as recently described [48]. To mimic biologically relevant conditions, flow cell assays were conducted using xylem sap extracted from 5-week-old Bonny Best tomatoes and sap was kept frozen at -20°C until use [80]. To synthesize the carboxymethyl cellulose-dopamine (CMC-DOPA) coating, an amidation reaction was induced between CMC carboxylic groups and the amine groups of DOPA. The resulting CMC-DOPA conjugate was alkalinized to facilitate bonding between the polymers and microfluidic channel surfaces. A 10 mg/mL CMC-DOPA solution was run through the system at the lowest flow rate of 10µL/h at 5 min intervals for 1 h. The systems were incubated overnight at 37°C, then washed with 3 mL of phosphate-buffered saline (PBS, pH 7.4) before use. Each system was seeded statically with a 10^9^ CFU/mL suspension of either GMI1000, *ΔlecF, ΔlecX*, *ΔlecF/X* or Δ*epsB* in xylem sap for 6 h to allow bacterial cells to attach to the channels. Then *ex vivo* Bonny Best xylem sap was pumped through the system at 38 µL/h by a microfluidic pump. The xylem sap was refreshed every 24 h and the experiment lasted 3 days. At the end of the experiment, biofilms were stained with 1% crystal violet, washed with sterile water three times, and visualized by a light microscope.

## Acknowledgements

We thank Dr. Sarah Swanson of the UW Newcomb Imaging Facility, José Sanchez- Gallego, and Stephanie Hayes for their technical support.

## References

1. Flemming HC, van Hullebusch ED, Neu TR, Nielsen PH, Seviour T, Stoodley P, et al. The biofilm matrix: multitasking in a shared space. Nat Rev Microbiol. 2023;21: 70–86. doi:10.1038/s41579-022-00791-0

2. Karygianni L, Ren Z, Koo H, Thurnheer T. Biofilm matrixome: Extracellular components in structured microbial communities. Trends Microbiol. 2020;28: 668–681. doi:10.1016/j.tim.2020.03.016

3. Flemming HC, Wingender J, Szewzyk U, Steinberg P, Rice SA, Kjelleberg S. Biofilms: An emergent form of bacterial life. Nat Rev Microbiol. 2016;14: 563–575. doi:10.1038/nrmicro.2016.94

4. Tsagkari E, Connelly S, Liu Z, McBride A, Sloan WT. The role of shear dynamics in biofilm formation. NPJ Biofilms Microbiomes. 2022;8: 33. doi:10.1038/s41522-022-00300-4

5. Sanfilippo JE, Lorestani A, Koch MD, Bratton BP, Siryaporn A, Stone HA, et al. Microfluidic-based transcriptomics reveal force-independent bacterial rheosensing. Nat Microbiol. 2019;4: 1274–1281. doi:10.1038/s41564-019-0455-0

6. Safni I, Cleenwerck I, De Vos P, Fegan M, Sly L, Kappler U. Polyphasic taxonomic revision of the *Ralstonia solanacearum* species complex. Int J Syst Evol Microbiol. 2014;64: 3087–3103. doi:10.1099/ijs.0.066712-0

7. Safni I, Subandiyah S, Fegan M. Ecology, epidemiology and disease management of Ralstonia syzygii in Indonesia. Front Microbiol. 2018;9: 419. doi:10.3389/fmicb.2018.00419

8. Yao J, Allen C. Chemotaxis is required for virulence and competitive fitness of the bacterial wilt pathogen *Ralstonia solanacearum*. J Bacteriol. 2006;188: 3697–3708. doi:10.1128/JB.188.10.3697-3708.2006

9. Yao J, Allen C. The plant pathogen *Ralstonia solanacearum* needs aerotaxis for normal biofilm formation and interactions with its tomato host. J Bacteriol. 2007;189: 6415–6424. doi:10.1128/JB.00398-07

10. Carter MD, Khokhani D, Allen C. Cell density-regulated adhesins contribute to early disease development and adhesion in *Ralstonia solanacearum*. Appl Environ Microbiol. 2023;89. doi:10.1128/aem.01565-22

11. Dalsing BL, Allen C. Nitrate assimilation contributes to *Ralstonia solanacearum* root attachment, stem colonization, and virulence. J Bacteriol. 2014;196: 949–960. doi:10.1128/JB.01378-13

12. Corral J, Sebastià P, Coll NS, Barbé J, Aranda J, Valls M. Twitching and Swimming Motility Play a Role in *Ralstonia solanacearum* Pathogenicity. mSphere. 2020;5: e00740–19. doi:10.1128/MSPHERE.00740-19/FORMAT/EPUB

13. Kang Y, Liu H, Genin S, Schell MA, Denny TP. *Ralstonia solanacearum* requires type 4 pili to adhere to multiple surfaces and for natural transformation and virulence. Mol Microbiol. 2002;46: 427–437.

14. Caldwell D, Kim B-S, Iyer-Pascuzzi AS. *Ralstonia solanacearum* differentially colonizes roots of resistant and susceptible tomato plants. Phytopathology. 2017;107: 528–536. doi:10.1094/PHYTO-09-16-0353-R

15. Rivera-Zuluaga K, Hiles R, Barua P, Caldwell D, Iyer-Pascuzzi AS. Getting to the root of *Ralstonia* invasion. Semin Cell Dev Biol. 2022;148–149: 3–12. doi:10.1016/j.semcdb.2022.12.002

16. Tsuzuki M, Inoue K, Kiba A, Ohnishi K, Kai K, Hikichi Y. Infection route in tomato roots and quorum sensing of *Ralstonia pseudosolanacearum* strain OE1-1. Physiol Mol Plant Pathol. 2023; 101995. doi:10.1016/j.pmpp.2023.101995

17. Lowe-Power TM, Khokhani D, Allen C. How *Ralstonia solanacearum* exploits and thrives in the flowing plant xylem environment. Trends Microbiol. 2018;26: 929–942. doi:10.1016/j.tim.2018.06.002

18. Arricau-Bouvery N, Duret S, Dubrana MP, Desqué D, Eveillard S, Brocard L, et al. Interactions between the flavescence dorée phytoplasma and its insect vector indicate lectin-type adhesion mediated by the adhesin VmpA. Sci Rep. 2021;11. doi:10.1038/s41598-021-90809-z

19. Vozza NF, Abdian PL, Russo DM, Mongiardini EJ, Lodeiro AR, Molin S, et al. A *Rhizobium leguminosarum* CHDL- (cadherin-Like-) lectin participates in assembly and remodeling of the biofilm matrix. Front Microbiol. 2016;7: 1608. doi:10.3389/fmicb.2016.01608

20. Rubeena AS, Abraham A, Aarif KM. Microbial lectins. Lectins: Innate immune defense and Therapeutics. Springer Nature; 2022. pp. 131–146. doi:10.1007/978-981-16-7462-4_7

21. Chemani C, Imberty A, De Bentzmann S, Pierre M, Wimmerová M, Guery BP, et al. Role of LecA and LecB Lectins in *Pseudomonas aeruginosa*-Induced Lung Injury and Effect of Carbohydrate Ligands. Infect Immun. 2009;77: 2065–2075. doi:10.1128/IAI.01204-08

22. Tielker D, Hacker S, Loris R, Strathmann M, Wingender J, Wilhelm S, et al. *Pseudomonas aeruginosa* lectin LecB is located in the outer membrane and is involved in biofilm formation. Microbiology (N Y). 2005;151: 1313–1323. doi:10.1099/mic.0.27701-0

23. Diggle S, Stacey R, Dodd C, Cámara M, Williams P, Winzer K. The galactophilic lectin, LecA, contributes to biofilm development in *Pseudomonas aeruginosa*. Environ Microbiol. 2006;8: 1095–1104. doi:10.1111/j.1462-2920.2006.01001.x

24. Passos Da Silva D, Matwichuk ML, Townsend DO, Reichhardt C, Lamba D, Wozniak DJ, et al. The *Pseudomonas aeruginosa* lectin LecB binds to the exopolysaccharide Psl and stabilizes the biofilm matrix. Nat Commun. 2019;10. doi:10.1038/s41467-019-10201-4

25. Pradhan BB, Ranjan M, Chatterjee S. XadM, a novel adhesin of *Xanthomonas oryzae* pv. *oryzae*, exhibits similarity to Rhs family proteins and is required for optimum attachment, biofilm formation, and virulence. Molecular Plant Microbe Interactions. 2012;25: 1157– 1170. doi:10.1094/MPMI-02-12-0049-R

26. Ghafoor A, Hay ID, Rehm BHA. Role of exopolysaccharides in *Pseudomonas aeruginosa* biofilm formation and architecture. Appl Environ Microbiol. 2011;77: 5238–5246. doi:10.1128/AEM.00637-11

27. Dharmapuri S, Sonti R V. A transposon insertion in the gumG homologue of *Xanthomonas oryzae* pv. *oryzae* causes loss of extracellular polysaccharide production and virulence . FEMS Microbiol Lett. 1999;179: 53–59. doi:10.1111/j.1574-6968.1999.tb08707.x

28. Tran TM, Macintyre A, Khokhani D, Hawes M, Allen C. Extracellular DNases of *Ralstonia solanacearum* modulate biofilms and facilitate bacterial wilt virulence. Environ Microbiol. 2016;18: 4103–4117. doi:10.1111/1462-2920.13446

29. Khokhani D, Lowe-Power TM, Tran TM, Allen C. A single regulator mediates strategic switching between attachment/spread and growth/virulence in the plant pathogen *Ralstonia solanacearum*. mBio. 2017;8: e00895–17. doi:10.1128/mBio.00895-17

30. Clough SJ, Flavier AB, Schell MA, Denny TP. Differential expression of virulence genes and motility in *Ralstonia (Pseudomonas) solanacearum* during exponential growth. Appl Environ Microbiol. 1997;63: 844–850. doi:10.1128/aem.63.3.844-850.1997

31. Kang Y, Saile E, Schell MA, Denny TP. Quantitative immunofluorescence of regulated eps gene expression in single cells of *Ralstonia solanacearum*. Appl Environ Microbiol. 1999;65: 2356–2362.

32. Kai K. The phc Quorum-Sensing System in *Ralstonia solanacearum* Species Complex. Annual Reviews. 2023;11: 213–231. doi:10.1146/annurev-micro-032521

33. Mcgarvey JA, Denny TP, Schell MA. Spatial-temporal and quantitative analysis of growth and EPS I production by *Ralstonia solanacearum* in resistant and susceptible tomato cultivars. Phytopathology. 1999;89: 1233–1239.

34. Denny TP, Baek S-R. Genetic evidence that extracellular polysaccharides is a virulence factor of *Pseudomonas solanacearum*. Molecular Plant-Microbe Interactions. 1991;4: 198–206.

35. Saile E, McGarvey JA, Schell MA, Denny TP. Role of extracellular polysaccharide and endoglucanase in root invasion and colonization of tomato plants by *Ralstonia solanacearum*. Phytopathology. 1997;87: 1264–1271. doi:10.1094/PHYTO.1997.87.12.1264

36. Milling A, Babujee L, Allen C. *Ralstonia solanacearum* extracellular polysaccharide is a specific elicitor of defense responses in wilt-resistant tomato plants. PLoS One. 2011;6: e15853. doi:10.1371/journal.pone.0015853

37. Kostlánová N, Mitchell EP, Lortat-Jacob H, Oscarson S, Lahmann M, Gilboa-Garber N, et al. The fucose-binding Lectin from *Ralstonia solanacearum*: A new type of β-propeller architecture formed by oligomerization and interacting with fucoside, fucosyllactose, and plant xyloglucan. Journal of Biological Chemistry. 2005;280: 27839–27849. doi:10.1074/jbc.M505184200

38. Sudakevitz D, Kostláno N, Blatman-Jan G, Mitchell EP, Lerrer B, Wimmerová M, et al. A new *Ralstonia solanacearum* high-affinity mannose-binding lectin RS-IIL structurally resembling the Pseudomonas aeruginosa fucose-specific lectin PA-IIL. Mol Microbiol. 2004;52: 691–700. doi:10.1111/j.1365-2958.2004.04020.x

39. Sudakevitz D, Imberty A, Gilboa-Garber N. Production, properties and specificity of a new bacterial L-fucose- and D-arabinose-binding lectin of the plant aggressive pathogen *Ralstonia solanacearum*, and its comparison to related plant and microbial lectins. J Biochem. 2002;132: 353–358. doi:10.1093/oxfordjournals.jbchem.a003230

40. Jacobs JM, Babujee L, Meng F, Milling A, Allen C. The in planta transcriptome of *Ralstonia solanacearum*: Conserved physiological and virulence strategies during bacterial wilt of tomato. mBio. 2012;3: e00114–12. doi:10.1128/mBio.00114-12

41. Meng F, Babujee L, Jacobs JM, Allen C. Comparative transcriptome analysis reveals cool virulence factors of *Ralstonia solanacearum* race 3 biovar 2. PLoS One. 2015;10: e0139090. doi:10.1371/journal.pone.0139090

42. Mori Y, Inoue K, Ikeda K, Nakayashiki H, Higashimoto C, Ohnishi K, et al. The vascular plant-pathogenic bacterium *Ralstonia solanacearum* produces biofilms required for its virulence on the surfaces of tomato cells adjacent to intercellular spaces. Mol Plant Pathol. 2016;17: 890–902. doi:10.1111/mpp.12335

43. McNeil M, Darvill AG, Fry SC, Albersheim P. Structure and function of the primary cell walls of plants. Ann Rev Biochem. 1984; 625–663.

44. Šulák O. Structure-function studies of lectins from opportunistic bacteria. 2009.

45. Orgambide G, Montrozier H, Servin P, Roussel J, Trigalet-Demery D, Trigalet A. High heterogeneity of the exopolysaccharides of *Pseudomonas solanacearum* strain GMI 1000 and the complete structure of the major polysaccharide. Journal of Biological Chemistry. 1991;266: 8312–8321.

46. Varbanets LD, Vasil’ev VN, Brovarskaya OS. Characterization of Lipopolysaccharides from *Ralstonia solanacearum*. Microbiology (N Y). 2003;72: 12–17.

47. Hamilton CD, Steidl OR, MacIntyre AM, Hendrich CG, Allen C. *Ralstonia solanacearum* depends on catabolism of myo-inositol, sucrose, and trehalose for virulence in an infection stage-dependent manner. Molecular Plant Microbe Interactions. 2021;34: 669–679. doi:10.1094/MPMI-10-20-0298-R

48. Chu LT, Laxman D, Abdelhamed J, Pirlo RK, Fan F, Wagner N, et al. Development of a tomato xylem-mimicking microfluidic system to study *Ralstonia pseudosolanacearum* biofilm formation. Front Bioeng Biotechnol. 2024;12. doi:10.3389/fbioe.2024.1395959

49. Sharma P, Johnson MA, Mazloom R, Allen C, Heath LS, Lowe-Power TM, et al. Meta- analysis of the *Ralstonia solanacearum* species complex (RSSC) based on comparative evolutionary genomics and reverse ecology. Microbal Genomics. 2022;8000791. doi:10.1099/mgen.0.000791

50. Hayashi K, Kai K, Mori Y, Ishikawa S, Ujita Y, Ohnishi K, et al. Contribution of a lectin, LecM, to the quorum sensing signaling pathway of *Ralstonia solanacearum* strain OE1-1. Mol Plant Pathol. 2019;20: 334–345. doi:10.1111/mpp.12757

51. Wilking JN, Angelini TE, Seminara A, Brenner MP, Weitz DA. Biofilms as complex fluids. MRS Bulletin. 2011. pp. 385–391. doi:10.1557/mrs.2011.71

52. Seminara A, Angelini TE, Wilking JN, Vlamakis H, Ebrahim S, Kolter R, et al. Osmotic spreading of Bacillus subtilis biofilms driven by an extracellular matrix. Proc Natl Acad Sci U S A. 2012;109: 1116–1121. doi:10.1073/pnas.1109261108/-/DCSupplemental

53. Mori Y, Hosoi Y, Ishikawa S, Hayashi K, Asai Y, Ohnishi H, et al. Ralfuranones contribute to mushroom-type biofilm formation by *Ralstonia solanacearum* strain OE1-1. Mol Plant Pathol. 2018;19: 975–985. doi:10.1111/mpp.12583

54. Tsumori C, Matsuo S, Murai Y, Kai K. Quorum Sensing-Dependent Invasion of Ralstonia solanacearum into *Fusarium oxysporum* Chlamydospores. Microbiol Spectr. 2023;11. doi:10.1128/spectrum.00036-23

55. Rusconi R, Stocker R. Microbes in flow. Curr Opin Microbiol. 2015;25: 1–8. doi:10.1016/j.mib.2015.03.003

56. Sauer MM, Jakob RP, Eras J, Baday S, Eriş D, Navarra G, et al. Catch-bond mechanism of the bacterial adhesin FimH. Nat Commun. 2016;7. doi:10.1038/ncomms10738

57. Thomas W. Catch bonds in adhesion. Annu Rev Biomed Eng. 2008;10: 39–57. doi:10.1146/annurev.bioeng.10.061807.160427

58. Audfray A, Claudinon J, Abounit S, Ruvoën-Clouet N, Larson G, Smith DF, et al. Fucose- binding lectin from opportunistic pathogen *Burkholderia ambifaria* binds to both plant and human oligosaccharidic epitopes. Journal of Biological Chemistry. 2012;287: 4335– 4347. doi:10.1074/jbc.M111.314831

59. Fujihashi M, Peapus DH, Kamiya N, Nagata Y, Miki K. Crystal structure of fucose- specific lectin from *Aleuria aurantia* binding ligands at three of its five sugar recognition sites. Biochemistry. 2003;42: 11093–11099. doi:10.1021/bi034983z

60. Bonnardel F, Kumar A, Wimmerova M, Lahmann M, Perez S, Varrot A, et al. Architecture and Evolution of Blade Assembly in β-propeller Lectins. Structure. 2019;27: 764–775.e3. doi:10.1016/j.str.2019.02.002

61. Mishra SK, Adam J, Wimmerová M, Koča J. In silico mutagenesis and docking study of *Ralstonia solanacearum* RSL lectin: Performance of docking software to predict saccharide binding. J Chem Inf Model. 2012;52: 1250–1261. doi:10.1021/ci200529n

62. Frensch M, Jäger C, Müller PF, Tadić A, Wilhelm I, Wehrum S, et al. Bacterial lectin BambL acts as a B cell superantigen. Cellular and Molecular Life Sciences. 2021;78: 8165–8186. doi:10.1007/s00018-021-04009-z

63. Wilhelm I, Levit-Zerdoun E, Jakob J, Villringer S, Frensch M, Übelhart R, et al. Carbohydrate-dependent B cell activation by fucose-binding bacterial lectins. Sci Signal. 2019;12.

64. Castro C, Ndukwe I, Heiss C, Black I, Ingel BM, Guevara M, et al. *Xylella fastidiosa* modulates exopolysaccharide polymer length and the dynamics of biofilm development with a β-1,4-endoglucanase. Vidaver AK, editor. mBio. 2023;14: 01395–23. doi:10.1128/mbio.01395-23

65. Gloag ES, Fabbri S, Wozniak DJ, Stoodley P. Biofilm mechanics: Implications in infection and survival. Biofilm. 2020;2: 100017. doi:10.1016/j.bioflm.2019.100017

66. Chew SC, Kundukad B, Seviour T, Van der Maarel JRC, Yang L, Rice SA, et al. Dynamic remodeling of microbial biofilms by functionally distinct exopolysaccharides. mBio. 2014;5: 1–11. doi:10.1128/mBio.01536-14

67. Wells M, Schneider R, Bhattarai B, Currie H, Chavez B, Christopher G, et al. Perspective: The viscoelastic properties of biofilm infections and mechanical interactions with phagocytic immune cells. Front Cell Infect Microbiol. 2023;13. doi:10.3389/fcimb.2023.1102199

68. Limoli DH, Jones CJ, Wozniak DJ. Bacterial Extracellular Polysaccharides in Biofilm Formation and Function. Microbiol Spectr. 2015;3. doi:10.1128/microbiolspec.mb-0011-2014

69. Ingel B, Caldwell D, Duong F, Parkinson DY, Mcculloh KA, Iyer-Pascuzzi AS, et al. Revisiting the Source of Wilt Symptoms: X-Ray Microcomputed Tomography Provides Direct Evidence That *Ralstonia* Biomass Clogs Xylem Vessels. PhytoFrontiers. 2022;2: 41–51. doi:10.1094/PHYTOFR-06-21-0041-R

70. Darsonval A, Darrasse A, Durand K, Bureau C, Cesbron S, Jacques MA. Adhesion and fitness in the bean phyllosphere and transmission to seed of *Xanthomonas fuscans* subsp. *fuscans*. Molecular Plant Microbe Interactions. 2009;22: 747–757. doi:10.1094/MPMI

71. Almeida RPP, Purcell AH. Patterns of *Xylella fastidiosa* colonization on the precibarium of sharpshooter vectors relative to transmission to plants. Ann Entomol Soc Am. 2006;99: 884–890.

72. Lowe-Power TM, Jacobs JM, Ailloud F, Fochs B, Prior P, Allen C. Degradation of the plant defense signal salicylic acid protects *Ralstonia solanacearum* from toxicity and enhances virulence on tobacco. mBio. 2016;7: e00656–16. doi:10.1128/MBIO.00656-16/FORMAT/EPUB

73. Monteiro F, Solé M, van Dijk I, Valls M. A chromosomal insertion toolbox for promoter probing, mutant complementation, and pathogenicity studies in *Ralstonia solanacearum*. Molecular Plant Microbe Interactions. 2012;25: 557–568. doi:10.1094/MPMI-07-11-0201/SUPPL_FILE/MPMI-07-11-0201E4.PDF

74. Perrier A, Barberis P, Genin S. Introduction of genetic material in ralstonia solanacearum through natural transformation and conjugation. Methods in Molecular Biology. Humana Press Inc.; 2018. pp. 201–207. doi:10.1007/978-1-4939-7604-1_16

75. Choi KH, Gaynor JB, White KG, Lopez C, Bosio CM, Karkhoff-Schweizer RAR, et al. A Tn7-based broad-range bacterial cloning and expression system. Nat Methods. 2005;2: 443–448. doi:10.1038/nmeth765

76. Lowe-Power T, Avalos J, Bai Y, Charco Munoz M, Chipman K, Tom CE, et al. A Meta- analysis of the known Global Distribution and Host Range of the *Ralstonia* Species Complex. bioRvix. 2022. doi:10.1101/2020.07.13.189936

77. Arkin AP, Cottingham RW, Henry CS, Harris NL, Stevens RL, Maslov S, et al. KBase: The United States department of energy systems biology knowledgebase. Nat Biotechnol. 2018;36: 566–569. doi:10.1038/nbt.4163

78. Letunic I, Bork P. Interactive Tree Of Life (iTOL) v5: an online tool for phylogenetic tree display and annotation. Nucleic Acids Res. 2021;49: W293–W296. doi:10.1093/nar/gkab301

79. Khokhani D, Tuan T, Lowe-Power T, Allen C. Plant Assays for Quantifying *Ralstonia solanacearum* Virulence. Bio Protoc. 2018;8: e3028. doi:10.21769/bioprotoc.3028

80. Georgoulis SJ, Shalvarjian KE, Helmann TC, Hamilton CD, Carlson HK, Deutschbauer AM, et al. Genome-Wide Identification of Tomato Xylem Sap Fitness Factors for Three Plant-Pathogenic *Ralstonia* Species. mSystems. 2021;6: e01229–21. doi:10.1128/msystems.01229-21

